# Using machine learning to predict quantitative phenotypes from protein and nucleic acid sequences

**DOI:** 10.1101/677328

**Authors:** David B. Sauer, Da-Neng Wang

## Abstract

**Background:** The link between protein or nucleic acid sequence and biochemical or organismal phenotype is essential for understanding the molecular mechanisms of evolution, reverse ecology, and designing proteins and genes with specific properties. However, it is difficult to practically make use of the relationship between sequence and phenotype due to the complex relationship between sequence and folding or activity.

**Results:** Here, we predict the originating species’ optimal growth temperatures of individual protein sequences using trained machine learning models. Both multilayer perceptron and k Nearest Neighbor regression outperformed linear regression could predict the originating species’ optimal growth temperature from protein sequences, achieving a root mean squared error of 3.6 °C. Similar machine learning models could predict organismal optimal growth pH and oxygen tolerance, and the quantitative properties of individual proteins or nucleic acids.

**Conclusions:** Using multilayer perceptron and k Nearest Neighbor regressions, we were able to build models specific to individual protein or nucleic acid families that can predict a variety of quantitative phenotypes. This methodology will be useful the *in silico* screening of individual mutations for particular properties, and also effective in the predicting the phenotypes of uncharacterized biological sequences and organisms.

## Background

The relationship between a protein or nucleic acid’s sequence and the biochemical or organismal phenotype is central to the study of evolution, reverse ecology, protein folding, and enzymatic activity. Further, knowing the relationship between a protein’s sequence and phenotype is also clinically and industrially valuable. For example, sequence differences (mutations) may reduce expression [1] or enzymatic activity [2], or cause the source organism to exhibit a disease phenotype [3]. Accordingly, highly tailored high-throughput experimental methods have been designed to efficiently screen the correspondence of sequence and phenotypes such as protein stability [4–6] and protein abundance [7]. Nevertheless, with the large number of sequence families and vast potential sequence space for each, more efficient computational methods are needed. However, the link between sequence and phenotype is often difficult to describe quantitatively due to the enormous potential sequence space, the complexities of protein and nucleic acid folding, and unknown biophysical and physiological mechanisms.

One sequence-phenotype relationship of particular interest is the connection between protein sequences and their thermodynamic stability. However, current *in silico* models of thermostability require a three-dimensional structure [8, 9], extensive mutagenesis of the target protein [10], or lack specificity to any particular protein family [11–14]. Fortunately, however, natural selection has already broadly sampled both sequence space and temperature space. Protein families often include homologs from organisms that grow at a wide range of temperatures. Organismal growth in each thermal niche places specific constraints on its proteins’ sequences such that the proteins are folded and functional under native conditions. Accordingly, studies comparing homologous proteins from species with different growth temperatures have identified sequence differences that correlate to the native thermal environment of the originating organisms [15–17]. Introducing corresponding mutations into model proteins often result in altered thermal stability or temperature-dependent activity, reflecting the role of these amino acids in thermoadaptation. Therefore, the available homologous protein sequences and organismal growth temperatures provide a large dataset for analyzing temperature-dependent protein properties, allowing novel analysis methods.

Here we report a quantitative method to predict the originating organism’s growth temperature (T_G_) from a protein sequence. The technique uses machine learning in k Nearest Neighbor (kNN), random forest (RF), and multilayer perceptron (MLP) models trained on an individual family of proteins or nucleic acids. Notably, no assumptions are made about chemical, structural, epistatic, or thermodynamic effects, and a three-dimensional structure is not required. Further, we show the protein’s predicted T_G_ is correlated with its experimentally determined melting temperature (T_M_) or temperature of optimal activity (T_A_). Thus, predicted organismal growth temperature 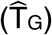 can serve as an easily calculable proxy for a protein’s thermal stability and temperature-dependent activity. Additionally, we demonstrate these regression methods’ generality by predicting other organismal phenotypes, and biochemical properties of individual protein and nucleic acid families.

### Construction of multilayer perceptrons

Setting out, we aimed to devise a method to predict quantitative phenotypes, such as T_G_, from a protein or nucleic acid sequence. As a part of making this a general method, we also want to avoid an explicit structure or description of the forces underlying protein or nucleic acid folding or organismal physiology. Therefore, we chose machine learning, which has been particularly useful when the relationship between the input and output is complex or unknown [44, 45]. Machine learning has been successfully applied to predicting a protein’s fold from sequence [46], the genotype of cancers from histopathology images [47], and the antimicrobial activity of a peptide sequence [48]. Similarly, here we applied machine learning in the form of multilayer perceptron (MLP), k Nearest Neighbor (kNN), and Random Forest (RF) regressions to quantitatively predict the originating organism’s growth temperature using protein sequences.

Generally, an MLP is a form of artificial neural network, a mathematical construct modeled on the structure and behavior of biological neural networks. As with a biological neural network, individual units (nodes or neurons) each accept and process input signals before producing an output (Figure S1A). In an MLP, these nodes are arranged into layers, termed “hidden layers”, with signals passed between consecutive layers, again mimicking the structure of biological neural networks. Starting from the input layer, each node’s value in the hidden layers is the result of an activation function applied to the weighted sum of the preceding layer’s nodes plus a layer bias value. The MLP output is then the weighted sum of the final hidden layer and an additional bias value.

The activation function of an MLP node is typically non-linear, mimicking the threshold potential and non-linear response of biological neurons. Central to its application here, MLPs with nodes that apply a non-linear activation function can act as universal approximators [49]. Therefore, we reasoned a sufficiently complex non-linear MLP could describe non-linear interactions, such as electrostatics and van der Waal’s contacts. Further, a non-linear MLP can model logical operators, such as AND and OR, and therefore could likely capture epistatic interactions. Consequently, we used MLPs with nodes that apply the non-linear, leaky rectified activation function (rMLPs) (Figure S1B).

As MLPs are mathematical models, the inputs are necessarily numerical. The inputs here are aligned sequences from a particular amino acid or nucleic acid family. We therefore convert the aligned sequences to sequences of Boolean variables (one-hot encoding) (Figure S2), where one or zero indicates the respective presence or absence of a particular amino acid or nucleic acid. We further removed positions from all one-hot encoded sequences that were absolutely conserved in the one-hot encoded training sequences, as these would not contribute to the regression. Therefore, one-hot encoding preserves the chemical sequence of a biopolymer in a numerical sequence of ones and zeroes. Notably, the one-hot encoding of the sequence does not contain a description of chemical or physical properties, minimizing the necessity of any assumptions of the relevant physical or chemical properties.

Regression methods, including machine learning, require the optimization of model parameters. Through “training”, the model weights and biases are iteratively refined using sequences with known regression target values, such as protein sequences and the associated originating organism’s optimal growth temperatures. However, it is essential to have mechanisms to avoid over-fitting and to independently evaluate accuracy [50]. Therefore, we used only 70% of the sequence-target pairs to optimize the MLP weight and bias parameters. The remaining 30% of the sequence-target pairs were excluded from training, and assigning to validation (10%) and test (20%) datasets. After each iteration of weight and bias optimization during training the validation sequences are predicted and compared to the known target values. Training is stopped when the validation Mean Square Error (MSE) no longer decreases to avoiding over-fitting of the model to features specific to the training dataset. Finally, the test dataset is used to calculate the MLP models’ accuracy, evaluating model generalizability and accuracy by predicting entirely unseen sequences. Notably, when placing sequences training, validation, or test datasets, identical sequences are placed in the same set. Therefore, the datasets have no sequences in common while retaining the same distribution as the input sequence alignment.

The MLP’s optimal number of nodes and their arrangement into layers - collectively the MLP’s “topology” - are not known *a priori*, and are likely specific to the protein or nucleic acid family and the target phenotype. Therefore, for each family, we considered all possible MLP topologies with the restrictions that the number of nodes in any hidden layer can range between two and twice the one-hot encoded sequence length, the network can have at-most 5 hidden layers, and the network must be over-determined. The number of possible topologies is vast, up to (2L - 1)^5^, where L is the one-hot encoded multiple sequence alignment length. Therefore, we applied an evolutionary algorithm to optimize the MLP topology [51], recombining and randomly permutating the lowest validation MSE topologies over multiple generations. Although this method does not evenly sample the entire topology space, and may not find the optimal topology, this method is empirically time efficient in finding an optimized MLP topology to predict phenotype.

We also explored the alternative machine learning methods of k-Nearest Neighbor and random forest regression. With kNN regression, we predicted the phenotype of a sequence as the average phenotype of the k closest training sequences, where distance is measured using the Hamming distance [52]. Notably, kNN regression uses only a single hyperparameter, k, which we optimized by minimizing the MSE versus the validation dataset. Random forest regression uses an ensemble of decision trees to predict the regression target, where each tree is a branching series of Boolean IF statements. RF regressions also have only a single hyperparameter, the number of trees in the forest, which we optimized by minimizing the MSE against the validation dataset. Finally, we again predicted the phenotype of test sequences to evaluate model accuracy.

## Results

### A trained MLP can predict organismal growth temperature from protein sequences

As an initial prototype for predicting a quantitative phenotype from a sequence, we examined the ability of an MLP to predict organismal growth temperature from protein sequences. Organismal growth temperature was selected as the regression target as T_G_s have been measured for many species [15]. Similarly, we expected proteins to be a useful input for such regression as many homologous protein sequences are known [26], and, most importantly, often contain adaptations to particular thermal niches [16, 17].

We first studied the Cold Shock Protein (CSP) family. The CSP family’s small protein size and strong conservation [53] results in many available CSP sequences relative to the protein length from species with a wide range of growth temperatures. This made the CSP family an ideal case study for regression of organismal growth temperature from protein sequence. Homologous cold shock protein sequences were collected from Pfam [26], extended by one amino acid at the C-terminus, and aligned in Promals3D [28]. In total, 41,972 homologous CSP sequences were identified with an available source organism growth temperature, with T_G_s measured -1.0 to 95.5 °C. All protein sequences were one-hot encoded, and MLPs were trained using the described algorithm. Of the 10^19^ possible topologies, using the evolutionary algorithm, 5,000 topologies were trained over 10 generations with 29,969 training sequences (71% of all sequences). The 3,956 validation sequences (9.4% of all sequences) were used to stop training and compare the various topologies’ accuracy. The trained MLP predicted the source organism growth temperature of 8,047 test sequences (19% of all sequences) with a root mean squared error of 4.0 °C (Pearson correlation r = 0.70). Notably, this MLP outperforms a linear regression trained with the same sequences (RMSE = 4.5 °C, r = 0.57), particularly in predicting the T_G_ of proteins from thermophiles.

In examining the trained MLP results, we noted the predicted T_G_ accuracy was poor for proteins from organisms with a growth temperature lower than 20 °C. This is perhaps due to these sequences’ rarity, as they comprise only 1.5% of the training species-T_G_ pairs. Additionally, training minimizes the squared error, which may lead to preferential optimization of sequences from thermophiles due to the T_G_ distribution’s positive skew. Finally, the mechanisms of adaptation from mesophilic to psychrophilic conditions may differ from the adaptations from thermophilic to mesophilic [54]. We therefore tested the accuracy of an MLP regression using only proteins from mesophiles and thermophiles. Excluding protein sequences from organisms with a growth temperature lower than 20 °C improved MLP accuracy (RMSE = 3.7 °C, r = 0.71). Therefore, we used only sequences from species with a T_G_ ≥20 °C in all subsequent growth temperature studies.

We further used the cold shock proteins from thermophiles and mesophiles of cold shock proteins to individually optimize the hyperparameters of MLP training, including the slope of the leaky ReLu function, the training batch size, and the learning rate. We found an optimal slope of 0.05, batch size of 500, and learning rate of 0.01 (Figure S3B-D). Training a MLP using these optimized hyperparameters resulted in a small increase in accurate when predicting the cold shock proteins’ originating species’ T_G_ (RMSE = 3.6 °C, r = 0.71) (Figure 1A), and outperformed a linear regression (Figure 1B). These optimized training hyperparameters were used throughout all subsequent studies.

**Figure 1.**
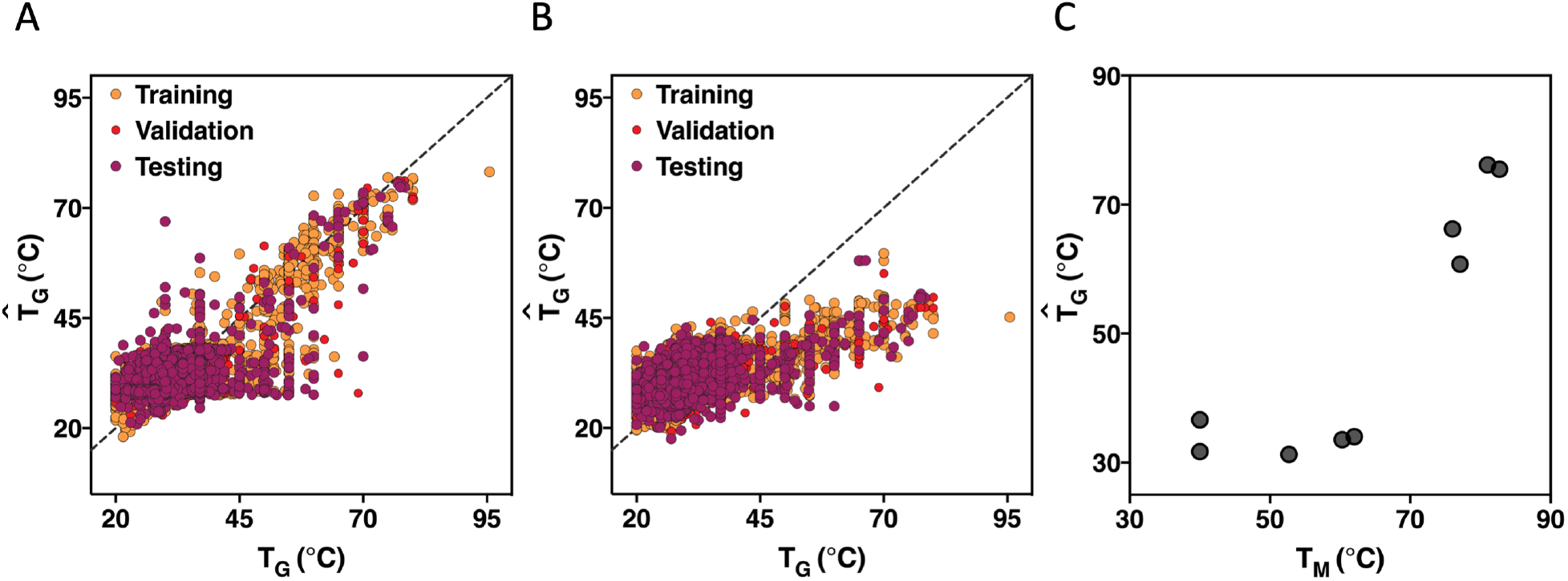
Organismal growth temperature can be predicted from the sequences of a protein family by using MLP regression. Regression of T_G_ using (A) the best rMLP or (B) linear regression model. The dotted line indicates perfect prediction. (C) Protein melting temperature versus predicted growth temperature of CSP homologs.

### Non-linearity is needed to predict growth temperatures

In examining the MLP topologies trained in the Cold Shock Protein regression, we found three distinct populations of model accuracy (Figure S3E). A population of topologies converged to low accuracy models (peak a). A second population is of topologies with accuracies similar to a linear regression (peak b). This set is unsurprising, as an rMLP can model a linear function. The final population of MLP models is more accurate than a linear regression (peak c), suggesting the rectified activation function is essential to regression accuracy.

However, as we trained multiple MLPs with many parameters, it was necessary to ensure that the improved accuracy of MLP regressions was not due to over-fitting or cherry-picking. Therefore, concurrent with the training of MLPs using a rectified activation function (rMLPs), we trained MLPs of the same topology with an identity activation function (iMLPs), where the activation function output is equal to the input. iMLPs are mathematically equivalent to linear regressions but fit the same number of parameters as MLPs with a rectified activation function for the same topology. As expected, these MLP regressions’ accuracy with an identity activation function is similar to the linear regression (Figure S3F). Notably, MLPs using rectified and identity activation functions have distinct distributions (Wilcoxon signed-rank test p < 10^−80^), confirming that the rectified activation function is essential to the improved prediction accuracy.

Notably, our experimental design always places identical sequences within the same training, test, or validation dataset. This strategy therefore reports the accuracy of a sequence sampled from a population with the same sequence space distribution as the initial dataset. This strategy also accounts for intra-sequence differences in T_G_ due to varying thermotolerance of proteins or organisms, or experimental errors in T_G_ measurement. However, this splitting strategy is less useful when describing accuracy for samples taken from a uniformly distributed sequence space, such as those generated during *in silico* screening. To examine this condition, we re-evaluated the regression accuracy for CSP sequences from thermophile mesophile using a test set where each unique sequence is present only once and associated with that sequence’s mean phenotype. Evaluated using this modified test set we found the rMLP model was still accurate (RMSE = 4.1 °C, r = 0.69), indicating that redundant sequences only modestly bias the accuracy metrics.

Finally, we performed ten repetitions of splitting the cold shock protein dataset and training rMLP regressions using different computational seed values. The consistent rMLP results (RMSE = 3.37 to 3.95 °C, r = 0.69 to 0.76) indicates that regression accuracy is not due to fortuitous sorting of the sequences or biases in the training.

### Predicted organismal growth temperature is correlated with experimentally determined melting temperatures of CSPs

We next explored if the rMLP predicted growth temperature would be of use in proteins’ biochemical characterization. Species’ T_G_ is known to correlate with its proteins’ thermal stability [55] and temperature-dependent activity [18]. We therefore examined if the predicted organismal growth temperatures of CSP proteins were similarly correlated with measured protein melting temperatures (Figure 1C). We found the homologs’ predicted growth temperatures and measured melting temperatures to be directly correlated (r = 0.88) [17, 56–69]. This correlation of 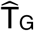 and T_M_ is similar to the correlation between measured T_G_ and T_M_ for the same sequences (r = 0.85).

It is worth noting the evolutionary selection of proteins protecting from cold shock complicates the theoretical relationship between T_G_ and T_M_. In principle, most proteins must be folded at the organismal growth temperature to support growth, imposing a theoretical bound on protein melting temperature such that T_G_ < T_M_. Broad analyses of many proteins from several organisms support this theory [55]. However, cold shock temperatures are necessarily lower than organismal growth temperatures. Consequently, the cold shock proteins’ melting temperatures must only be greater than the cold shock temperatures to support organismal growth. Therefore, the melting temperatures of cold shock proteins could in principle be lower than the originating organism’s growth temperature. However, we observe the homolog’s T_M_s are still greater than both rMLP predicted T_G_s and measured T_G_s of each of the originating species. This may indicate the temperature difference between optimal organismal growth and the physiological onset of CSP activity is generally small, or there are other functions of CSP homologs at the organisms’ optimal growth temperatures [70].

### Adenosine kinases’ biochemical characteristics correlate with 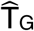

We next further examined if the rMLP predicted organismal growth temperature correlated with the protein’s stability or activity. We therefore applied the same method to Adenosine Kinases (ADK), an extensively studied [16, 71, 72] family that catalyzes the interconversion of adenosine nucleotides. As observed in other proteins [18], we found that the ADK homologs’ temperatures of optimal activity [16, 73] were correlated with measured originating species’ growth temperatures (r = 0.79). While there were too few ADK sequences to train an over-determined MLP, a linear regression was already highly accurate at predicting the originating species’ growth temperature (RMSE = 4.0 °C, r = 0.80) (Figure 2A). We found the 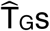 for characterized ADK homologs and reconstructed ancestral sequences correlated with protein T_M_ (r=0.72) (Figure 2B) and T_A_ (r = 0.61) (Figure 2C), indicating that 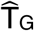 could serve as a proxy for T_A_ and T_M_.

**Figure 2.**
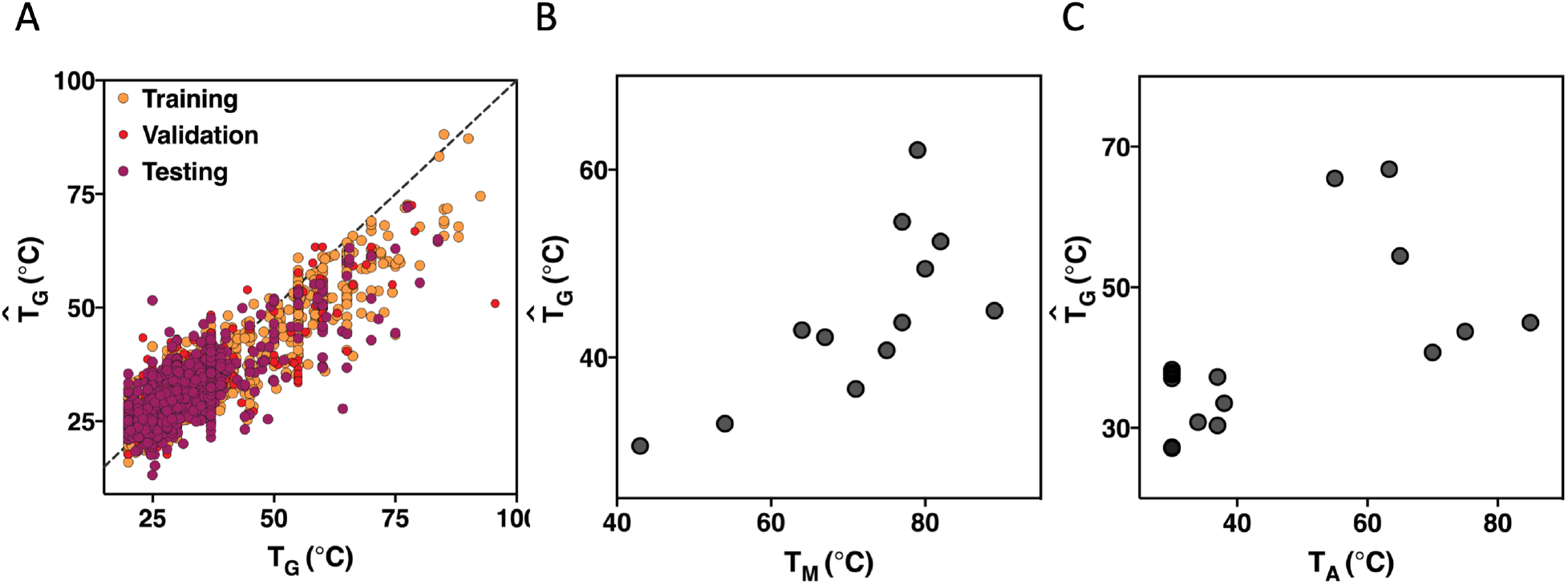
Linear regression predicted T_G_s correlate with ADK’s biochemical properties. (A) Linear regression of T_G_ from ADK sequences. The dotted line indicates perfect prediction. Predicted growth temperature versus (B) protein melting temperature and (C) temperature of optimal activity of ADK proteins.

### Thermophilic proteins have unique adaptations to temperature

We noted that 97% of the CSP sequences available are from mesophiles (Figure S4A). This was unsurprising given the bias of the characterized organisms [15, 18]. While random sampling ensures the train, validation, and test datasets have the same distribution as the input alignment, it was necessary to explore if the non-uniform distribution in phenotypes affected rMLP accuracy.

We first considered if sequences from mesophiles alone sufficiently sampled sequence space to accurately predict the T_G_ of homologs from thermophiles (Figure S4B). If successful, this would indicate that the thermoadaptive sequence differences between homologs from mesophiles and thermophiles are contained within the sequence space sampled by mesophiles alone. However, limiting the training and validation datasets to only proteins from mesophiles reduced regression accuracy (RMSE = 4.2 °C, r = 0.57), with a systematic under-prediction of proteins from thermophiles. Therefore, proteins from thermophiles likely contain amino acids at particular positions outside the sequence variation seen within CSPs from mesophiles.

We also investigated if the non-uniform distribution of organismal growth temperatures in the training dataset hindered the regression’s accuracy. This would be possible if, during training, the optimization of MLP weights and biases was dominated by the numerous protein sequences from mesophiles. While we expect this risk to be minimized by using mean square error when optimizing rMLPs, rather than mean absolute error, we also tested a “balancing” strategy for mitigating the effects of sampling non-uniformity. In balancing, we generated a new cold shock protein training dataset where rare thermophile sequences are over-sampled to be equal in proportion to the more common mesophiles (Figure S4C). The validation and test datasets remained unchanged. Applying this new training dataset, the accuracy of the rMLPs in predicting the unseen test dataset was worse than without balancing (RMSE = 4.4 °C, r = 0.65) (Figure S4D). As the number of unique CSP sequences from thermophiles is much smaller than those from mesophiles, in this case the oversampling of sequences from thermophiles may have led to over-fitting of inconsequential amino acids unique to these sequences.

Together, these results make clear that the relatively few (3%) sequences from thermophiles in the training dataset are necessary and sufficient for the prediction of optimal growth temperature of homologs from thermophiles. The non-uniform distribution of protein sequences and species T_G_s does not appear to have harmed regression accuracy, though accuracy may increase with more unique homologs from thermophiles having associated organismal growth temperatures.

### rMLPs are more accurate than linear regressions even with relatively few sequences

The ability of an rMLP to model increasingly complex functions is dependent upon increased network depth and width. However, as network topology is required to be over-determined, network complexity is limited by the number of training sequence – organismal growth temperature pairs. To examine how regression accuracy scales with the number of sequences, we generated smaller CSP training and validation datasets by random sampling. The test set remained unchanged, in order to consistently measure regression accuracy. While it was not possible to build over-determined MLPs with 10% of the training sequences, the rectified MLP function outperformed an identity activation function with only 20% of the training and validation sequences (Figure 3). This suggests that as few as 3.9 training sequences per one-hot encoded amino acid, or 20.2 sequences per column of the multiple sequence alignment, are sufficient to capture non-linear effects on the relationship between protein sequence and organismal optimal growth temperature.

**Figure 3.**
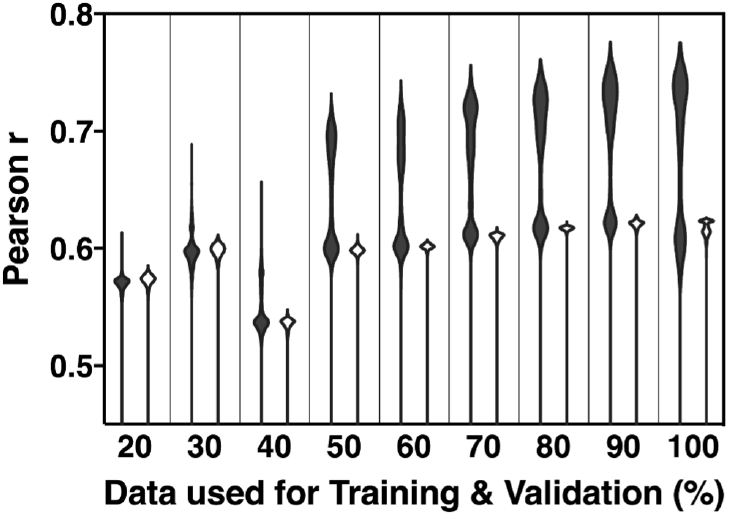
The proportion of non-linear MLP topologies outperforming equivalent topologies with a linear activation function increases with more training data. MLP accuracy trained using subsets of the training and validation sequences with either rectified (filled) or identity (unfilled) activation functions.

### Particular amino acids are critical to T_G_ prediction

When exploring possible MLP topologies, we only considered those over-determined topologies, with more training sequences than model parameters. As the number of model parameters depends on the one-hot encoded sequence length, we realized that longer or less well-conserved protein families might be precluded from this type of analysis. Fortunately, previous studies had indicated that only a small fraction of mutations to a protein’s sequence alter protein stability [5, 6, 17]. We therefore hypothesized that most sequence differences were neutral to thermoadaptation, analogous to passenger mutations. Thus, most one-hot encoded amino acids would not contribute to the regression’s accuracy, while potentially adding noise to the regression and decreasing the maximum complexity of the topologies examined. To test this hypothesis, we examined the correlation of each one-hot encoded position with T_G_ and whether excluding un-correlated amino acids would improve regression accuracy.

We identified the first-order correlation between amino acid presence or absence and the originating species’ growth temperature using the point-biserial correlation coefficient (Figure S5A). Using only the encoded amino acids with the greatest absolute correlation causes a modest loss in accuracy (RMSE = 3.7 °C, r = 0.70) while still outperforming a linear regression (RMSE = 4.7 °C, r = 0.46) using only 10% of the encoded protein sequence (Figure S5B). We similarly used a fit top-hat function to identify amino acids with a second-order correlation to growth temperature (Figure S5C). Again using only 10% of the encoded amino acids, those with the greatest absolute correlation to a top-hat function, an rMLP could predict growth temperature with a root mean squared error of 3.9 °C (r = 0.66) (Figure S5D).

These results confirm that only a subset of amino acids in the CSP family’s alignment are correlated with temperature and needed to predict the originating species’ growth temperature. Therefore, using a similar point-biserial or top-hat correlation threshold would increase the effective data-to-parameter ratio, and allow for the over-determined regression of longer proteins. Alternatively, deeper and broader topologies could be used for shorter proteins, improving accuracy by accounting for more complex interactions in the sequence.

### Pruning under-determined MLPs improve prediction accuracy

While limiting the regression inputs clearly can produce rMLPs as accurate as using all sequences, we recognized that applying a first or second-order threshold would necessarily remove amino acids with weaker or more complex correlations to the phenotype. To better balance MLP complexity and the data-to-parameter ratio, we explored model pruning or the selective removal of weakly contributing model weights. In this way, an under-determined model can initially be trained using all amino acids, and more nodes per layer can be explored, with the final weights pruned such that the model is ultimately over-determined.

We trained rMLP models of CSP sequence to the originating species’ T_G_ as previously, allowing the MLP topology to be up to 10-fold underdetermined. We then scanned weight sparsities, finding up to 80% of the model weights could be set to zero without an appreciable loss of accuracy. Calculating each pruned model’s data-to-parameter ratio, there is a notable region of sparsity between 71 to 80% where the model is over-determined without a loss of accuracy. Pruning an initially under-determined model to 71% sparsity, the resulting pruned rMLP model could predict growth temperature with a root mean squared error of 3.6 °C (r = 0.72) (Figure S5E). Therefore, while pruning did not improve accuracy in this case, this method offers the same benefits as the point-biserial or top-hat thresholds without making assumptions *a priori* of the most critical amino acids to regression accuracy.

### Non-linear MLPs in the regression of other protein families

Next, we examined if rMLPs could be used as a general method for predicting organismal growth temperature. Therefore, we trained MLP regression models to predict the originating species’ T_G_ for other protein families (Table S1). These families included interaction and enzymatic domains, targeted to various cellular localizations, and alpha-helical and beta-barrel membrane proteins. Of the 50 protein families examined, 38 had a sufficient number of sequences to train an over-determined MLP. The rMLP predicted growth temperature was correlated with measured growth temperature for all 38 families, and in all but two cases outperformed a linear regression. Species growth temperature predictions were consistent across all rMLP regression (mean species RMSD = 2.4 °C), with a mean species RMSE = 4.0 °C.

### MLPs can predict biochemical phenotypes of proteins and nucleic acids

With the success of rMLPs to predict organismal growth temperature from various protein families, we hypothesized that the same method could be applied to regression of other quantitative phenotypic traits from the aligned sequences of other biopolymers.

We first examined the ability of MLPs to predict the phenotype of densely sampled sequence space by calculating regression on deeply mutagenized sequences of the human Yes Associated Protein 65 WW domain [32] and their corresponding binding affinities (Figure 4A). The training rMLP’s predictions were strongly correlated with measured affinities (r = 0.87). Similar rMLP regressions could predict brightness for the fluorescent proteins eqFP611 (r = 0.96) and avGFP (r = 0.97) [34, 37] (Figure 4B and Figure S6A).

**Figure 4.**
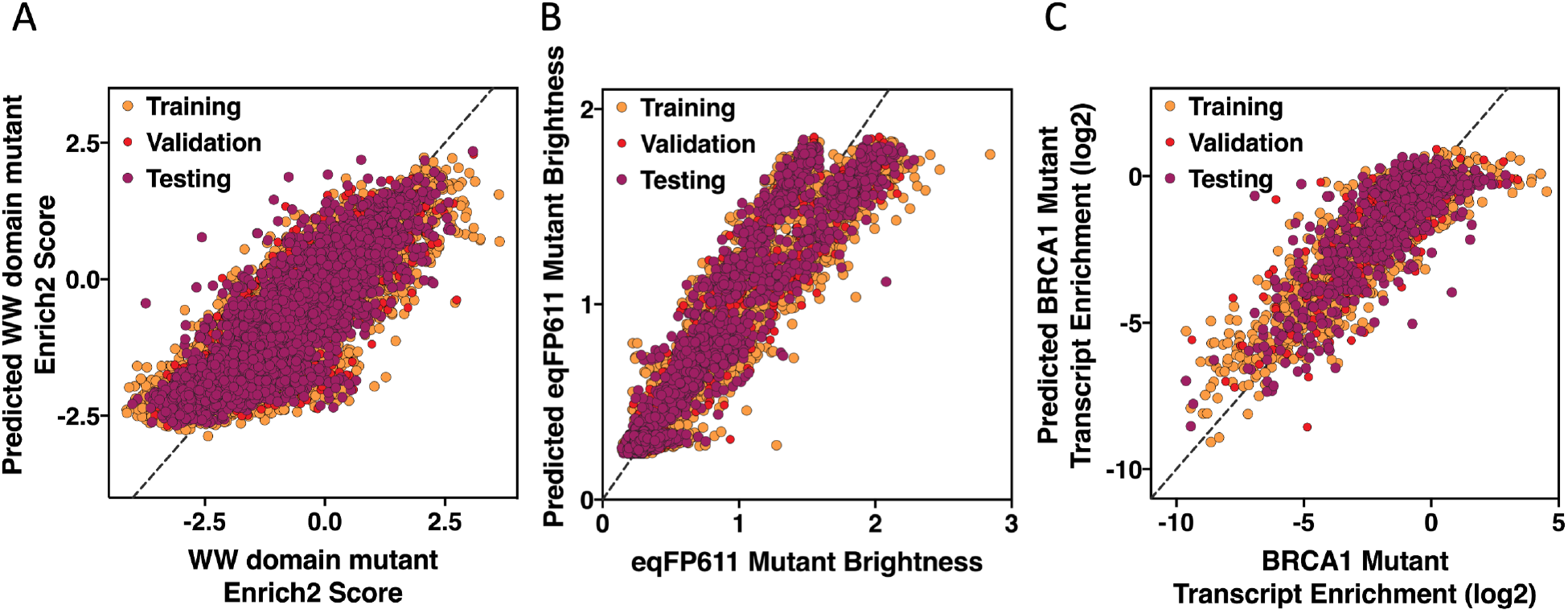
Non-linear MLPs can predict other phenotypes from protein and nucleic acid sequences. Predicting the (A) enrichment of WW domain mutants and (B) brightness of eqFP611 mutants and (C) transcript enrichment of mutant BRCA1 nucleic acid sequences using trained rMLP models. The dotted lines indicate perfect prediction.

Noting that our one-hot encoding makes no assumptions of the physical or chemical characteristics of the biopolymer’s monomers, we next tested if nucleic acids can be analyzed by the same method. We therefore applied the regression method to mutant sequences of the BRCA1 gene [36], finding the rMLP’s predictions strongly correlated with measured transcript enrichment (r = 0.83) (Figure 4C). Additionally, rMLPs predicted the catalytic activity of an engineered guanine-inhibited ribozyme both with (r = 0.89) and without (r = 0.90) ligand [35] (Figure S6B&C).

These results verify this method’s generality to predict various quantitative and qualitative phenotypes from proteins and nucleic acid sequences.

### The contribution of sequence identity to MLP accuracy varies by protein family

Describing the relationship between inputs and outputs of machine learning methods is often challenging, with a trained model treated as a “black box”. However, this can limit the interpretability of the produced models and their applicability to particular problems. Notably relevant here is the possibility that the MLPs’ predictions are based upon sequence similarity, rather than the expected biophysical interactions of the amino acids. This is a particular of naturally-evolved biomolecules, where shared phylogeny and biases in the organisms’ sequenced may result in many highly related sequences.

To evaluate the ability of considering sequence similarity alone in predicting phenotypes, we calculated k-Nearest Neighbor regression (kNN) of the same sequences, a method that explicitly applies similarity to predict a target value. In a kNN regression, an unknown protein sequence’s predicted phenotype is simply the average phenotype of the k most closely related sequences from the training set. Optimizing k using the validation dataset, we found that a kNN predicted CSP T_G_s with an accuracy of 3.5 °C (r = 0.73) (Figure S7A). Over all the sequence families analyzed, kNNs are more accurate in most naturally evolved sequences, while MLPs outperform kNN regressions with the many of the deeply mutagenized sequences. Notably, the difference in these methods’ performance did not appear to correlate with the fraction of sequence space sampled (Figure S7B). Similarly, the difference in performance between MLP and kNN regressions did not correlate with the number of training sequences per one-hot monomer (Figure S7C), suggesting that sequencing depth does not underly this difference. We therefore hypothesize that the performance difference of MLP and kNN regressions is due to the unbiased sampling of sequence space in deep mutagenesis experiments. We would therefore expect MLP’s relative performance on naturally evolved sequences to improve as more diverse organisms are sequenced.

Going one step further, we wanted to compare the rMLP and kNN results to describe the contribution of sequence identity to rMLP regressions. However, this is difficult as both methods are optimized to predict the same value. Fortunately, if rMLP models predicted on sequence identity, as kNN models operate, we would expect the residual errors in the predictions to be highly correlated. We therefore compared the test prediction errors of both methods in all regressions. Over all the regressions, the residual kNN error was only moderately correlated with rMLP residual error for naturally evolved (mean r = 0.68) and deep mutagenesis sequences (mean r = 0.58). Nevertheless, with this difference in error despite predicting the same test sequences, we conclude that sequence identity is a component of rMLP accuracy which varies between the individual sequence family regressions.

We also explored Random Forest (RF) regressions to predict phenotypes from sequences. As with rMLPs, this method is sensitive to differences of individual amino acids or nucleotides. However, while rMLPs capture the epistatic interactions through the weights between layers, in RF regressions the trees must sequentially discriminate on each interacting residue. While it is enticing to consider studying the Boolean logic of individual decision trees responsible for phenotype determination, the number trees within a random forest makes this impractical. Nevertheless, we were curious if random forest regressions could effectively prediction phenotypes from naturally and synthetically evolved sequences. Optimizing the number of trees in each random forest regression using the validation dataset, we found that RF regressions were the most accurate machine learning method of predicting phenotype for a subset of naturally evolved sequences. However, rMLPs clearly outperform fandom forests in predicting synthetically generated sequences. These results suggest while decision trees are useful for predicting phenotype from naturally evolved sequences, this method may not efficiently or adequately describe the general phenotype-sequence relationship of more deeply or uniformly sampled sequence space.

## Discussion

The identification or design of biopolymers with particular biochemical or biophysical properties is often central to their study or their use in industrial applications. However, the link between sequence and phenotype is often challenging to describe, due to the large potential sequence space and the difficulty of characterizing individual protein sequences.

Here, we successfully generated mathematical models to predict the various quantitative phenotypes from protein and nucleic acid sequences. Growth temperatures could be predicted from protein sequences with a root mean squared error of 3.6 °C. The predicted T_G_s correlate with experimentally determined melting temperatures and temperatures of optimal activity. Similarly, various biochemical activities could be predicted from biopolymers’ sequences with strong correlation to the measured phenotypes. As phenotypically characterizing an individual sequence is typically tedious and time-consuming, these regression methods of predicting phenotype *in silico* from sequence offer significant advantages over experimental techniques.

### Linear regressions are sufficient for some protein families

Some phenotypes can clearly be modeled as the linear combination of individual amino acid contributions, such as thermostability as seen in some membrane [74] and soluble proteins [17], and prediction of T_G_ from ADK protein sequences. However, non-linear effects are seen in the thermal stability of the Arc repressor [75], the brightness of the eqFP611 fluorescent protein [34]. Similarly, non-linear effects are seen here in the regression of organismal growth temperature from sequences. The inconsistent success of linear regression models in predicting organismal growth temperature from sequence supports the hypothesis that the physical interactions that underlie thermoadaptation vary by protein family [76].

As multilayer perceptrons are universal approximators this algorithm likely represents a general solution to describing the relationship between sequence and quantitative characteristics of the protein. However, our results also highlight the value of k nearest neighbor and random forest regressions when analyzing naturally evolved sequences, likely due to shared phylogeny or uneven sampling of sequence space.

### Predicting single mutant’s effects requires densely sampled sequence space

In principle, a trained rMLP should be sensitive to the effects of a single or few amino acid differences, such as experimentally generated point mutations. We therefore examined the correlation of rMLP predicted growth temperatures to the measured melting temperatures for single and double mutants of a CSP ortholog from *Bacillus subtillis* (BsCSP) (Figure S8A). We found no correlation between mutant protein melting temperatures [66, 68] and predicted organismal optimal growth temperatures calculated from the mutant proteins’ sequences (r = 0.17). Comparing BsCSP to the training sequences, we noted that homologs with high sequence identity to BsCSP come from organisms with T_G_s similar to *Bacillus subtillis* (Figure S8B). This contrasts with BsCSP mutants, with only one or two amino acid changes, exhibiting significantly altered melting temperature from wild type [17, 66, 68]. We therefore suspect that the available CSP sequences do not sufficiently sample sequence space to capture the effects of few or rare sequence differences. This hypothesis is supported by the accuracy of rMLP in predicting the binding affinity of deeply mutagenized sequences. In particular, the WW domain dataset consists of only single, double, and triple mutants, corresponding to 91-97% sequence identity. Therefore, this result demonstrates that rMLPs can only accurately predict the effects of a few mutations with sufficiently sampled sequence space.

### Predicting other organismal phenotypes

While other methods can predict organismal growth temperatures [14, 77], we noted that rMLP regression can calculate 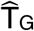 without requiring a complete genome or proteome sequence for the organism. This is particularly useful if the organism of interest no longer exists, such as ancestral organisms, predicting their thermal niche by analyzing their reconstructed sequences. Though the thermal niches of ancestral organisms have been inferred by the experimental characterization of reconstructed proteins’ melting temperatures [16] predicting organismal growth temperature *in silico* regression models are faster. Furthermore, growth temperature only places a minimum on protein melting temperatures (T_M_ > T_G_) and a rough bound on enzymes’ temperature of optimal activity (T_A_ ≈ T_G_ for enzymes limiting the growth rate). Therefore, predicting T_G_ from sequence may also be more accurate than biochemically characterizing individual proteins. With this success in predicting T_G_, we next explored if similar regression could predict other organismal phenotypes from sequences and demonstrate a general reverse ecology tool. We therefore calculated regressions for the oxygen tolerance and optimal growth pH of microorganisms.

Regressions predicting the originating species’ optimal growth pH from protein sequences are challenged by the relatively few 4822 species with reported growth pH. Limiting regressions to short, highly-conserved domains, we were still unable to calculate overdetermined regressions. We therefore calculated only linear, kNN, and RF regressions for the periplasmic YceI protein [78] and the C-terminal domain of 3-hydroxyacyl-CoA dehydrogenase [79]. Using random forest regressions, we could predict species optimal growth pHs with an RMSE of 0.64 and 0.71 for each family, respectively (Figure 5A).

**Figure 5.**
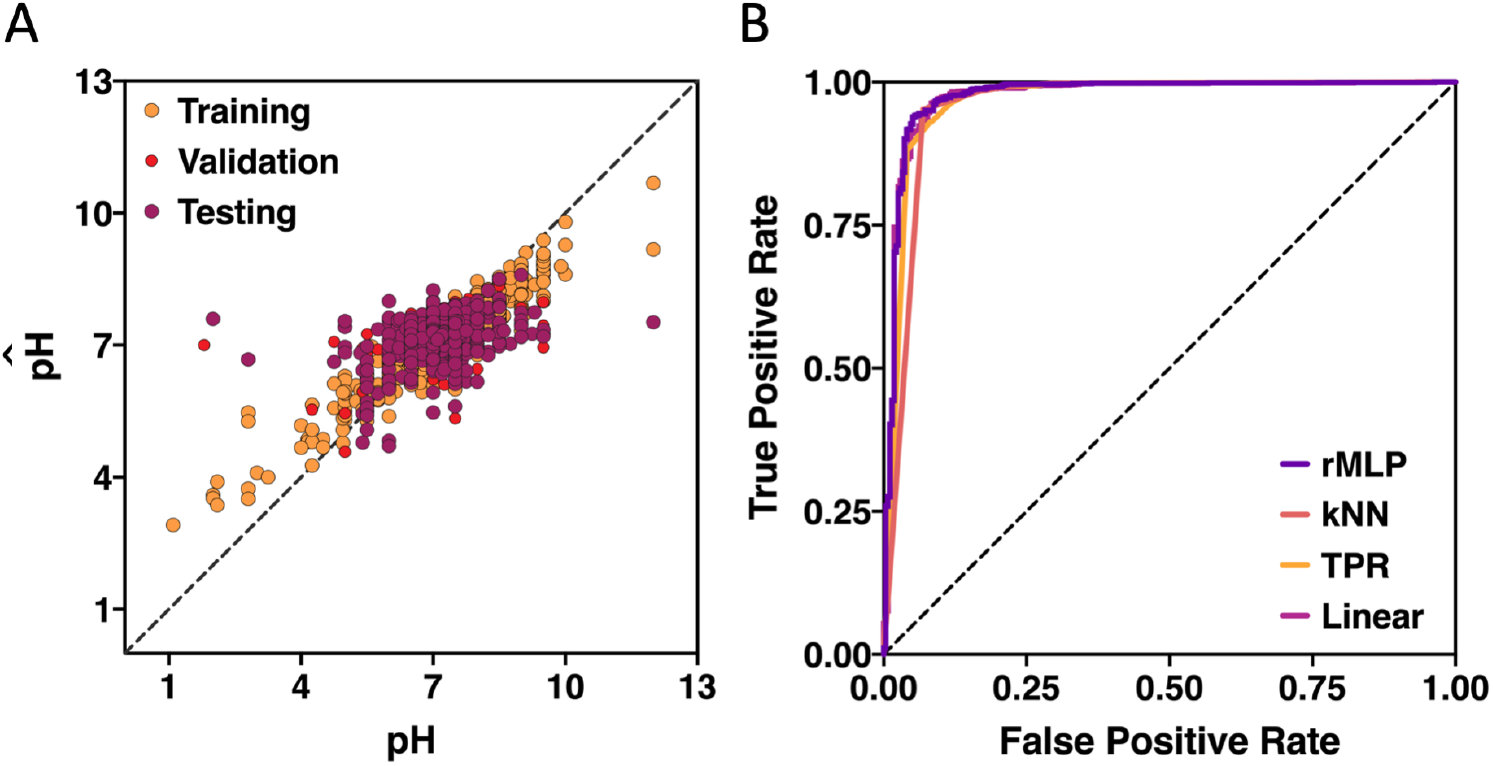
Machine learning can predict other organismal phenotypes from protein sequences. (A) Predicting the pH from 3-hydroxyacyl-CoA dehydrogenase’s C-terminal domain using a random forest regression. (B) ROC curves of rMLP, kNN, RF and linear regressions of oxygen tolerance from Fe/MnSOD N-terminal domain sequences. The dotted line indicates the line of non-discrimination.

In contrast, while oxygen tolerance is more widely reported for many microorganisms, it is usually reported as a categorical value of aerobic or anaerobic though in reality this is a continuous quantitative trait. However, we kept with convention and numerically encoded the organismal oxygen tolerance as 1 for aerobic and 0 for anaerobic for the 7238 characterized microbes. We excluded species with intermediate or poorly defined oxygen tolerance, such as microaerophiles, facultative anaerobes, and aerotolerant anaerobes. We then calculated regressions as previously for the N- and C-terminal domains of Iron/Manganese Super Oxide Dismutase (Fe/MnSOD), a protein essential for oxygen tolerance [80]. We found a rMLP regression was the most accurate in classifying the source organism’s oxygen tolerance using the N-terminal domain sequences, with an AUC of 0.98. In contrast a random forest was most accurate when evaluating C-terminal domain sequences (AUC = 0.98) (Figure 5B).

Collectively, these results indicate regressions can quantitatively predicting an organism’s ecological niche from its sequences. Notably, the CSP, 3-hydroxyacyl-CoA dehydrogenase, and Fe/MnSOD proteins are all highly conserved and therefore may prove useful in the routine reverse ecology characterization of ancestral or uncharacterized organisms’ T_G_, optimal growth pH, and oxygen tolerance.

## Conclusions

We demonstrate the use of machine learning models to predict quantitative phenotypes from the sequences of proteins and nucleic acids. This protocol makes no assumptions about the input data (i.e. it uses only the provided sequences), and thus requires significantly less prior knowledge than other predictors. Additionally, these *in silico* methods typically take a few hours to complete on a personal computer and therefore are significantly faster than experimental determinations of phenotype. Finally, as this machine learning strategy can predict organismal phenotypes from biological sequences, our methodology represents a novel reverse ecology strategy for describing uncharacterized and ancestral organisms.

## Methods

### Sequence and organismal growth temperature collection

Species’ measured growth temperatures were collected from published sources [15, 18–22] and the BacDive [23], Riken BRC [24], BCCM [25] cell line and strain catalogs. Species’ measured optimal growth pHs were downloaded from BacDive. Duplicate values of the same species were averaged. Species’ oxygen tolerances were downloaded from BacDive, with the statistical mode used when multiple values were recorded. Species annotated as facultative anaerobes, facultative anaerobes, and microaerophiles were excluded from further analysis. Oxygen tolerance was then numerically encoded as 1 for aerobes and 0 for anaerobes.

Protein domain sequence alignments were downloaded from the Pfam 33.1 database [26] and used without modification unless otherwise noted. Reconstructed ancestral Adenosine Kinase (ADK) sequences [16] were combined with extant proteins from the Pfam alignment. Cold Shock Protein (CSP) sequences identified in Pfam were extended at the C-terminus by one amino acid using the sequences in UniProtKB release 2019_08 [27]. CSP and ADK sequences were then re-aligned in Promals3D [28]. Proteins inducing gaps, annotated as fragments, or with residues outside the standard protein alphabet (ACDEFGHIKLMNPQRSTVWY-) were excluded.

### Data division into training, validation, and test datasets

The method requires three datasets for training the regression model (training), preventing overfitting to the training dataset (validation), and final evaluation of model accuracy (test). Therefore, identical sequences were clustered, and these clusters were then randomly assigned into training (70%), validation (10%), or test (20%) datasets.

Fractional sequence space was calculated as U/N^L^ where U is the number of unique sequences, L is the number of non-absolutely conserved positions in the alignment, and N is the number of possible monomers for that biopolymer type (5 for DNA or RNA, and 21 for proteins). Sequence families with sampling of sequence space so small as to call a number error were excluded from the comparison of sequence space sampling and regression accuracy.

### Sequence assignment, encoding, and balancing

For T_G_ regression, species assignments for each protein were collected from UniProtKB release 2019_08. Those sequences without a species assignment or an originating species’ T_G_ were excluded from further analysis. All CSP and ADK sequences with a characterized T_A_ and T_M_ were removed from the alignments; and used only to compare T_G_ and 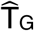 to T_M_ and T_A_. Regression of oxygen tolerance used the same procedure, replacing T_G_ with the originating species’ numerically encoded oxygen tolerance.

All sequences were then one-hot encoded, with invariant positions of the one-hot training sequences removed from all alignments.

The training dataset was balanced by first calculating a histogram of training sequence phenotypes with 20 bins. In addition to all the sequences in the original alignment, sequences were added to the alignment by random selection with replacement from each bin until all bins had the same number of sequences as the most populous bin.

Phenotype-correlated positions were identified using the point-biserial correlation or a top-hat function fit to the relationship between phenotype and the one-hot encoded amino acid. When thresholded, the positions were ranked by absolute top-hat function Pearson correlation or absolute point-biserial correlation, and only the top 10% were used for regression.

### MLP training

The multilayer perceptrons (MLPs) were trained using the identity or leaky ReLu activation functions [30]. All regressions were trained using the Adam solver [31], with the mean square error (MSE) as the loss function. Model checkpoints were saved at each epoch if the current model validation dataset MSE was lower than the previous checkpoint. The training was stopped when the validation dataset MSE did not decrease for two consecutive training epochs.

### Topology generation and search

MLP architectures were built systematically, requiring only that the first hidden layer have at most twice as many nodes as the input layer. Topologies were limited to between 1 and 5 hidden layers, in addition to the input and output layers. Of the potential topologies, 500 were randomly selected and trained per generation for 10 generations. After each generation, the top scoring 20% of the topologies (based on the MSE of the validation dataset) were recombined and mutated, and used as input for the following generation. Recombining topologies consisted of joining two topologies at a random layer chosen from each. Topologies were mutated by randomly changing the number of nodes in a randomly chosen layer. Finally, the Pearson correlation coefficient, mean squared error, and root mean square error was calculated and reported using the test dataset for the lowest validation MSE model.

### MLP hyperparameter tuning

The initial MLP hyperparameters included a leaky ReLu slope of 0.01, batch size of 500, and learning rate of 0.1. After the initial demonstration of MLP regressions on psychrophiles-mesophiles-thermophiles and mesophiles-thermophiles datasets, hyperparameters were tuned individually using the thermophiles-mesophiles dataset. Leaky ReLu slope optimization screened values of 0, 0.001, 0.005, 0.01, 0.05, 0.1 and 0.5. Batch sizes tested were 100, 500, 1000, 5000, and 10,000, and learning rates of 0.001, 0.01, 0.1, and 1 were also evaluated. The individual hyperparameters which gave the minimum MSE of the test set were selected as optimal.

### MLP pruning

MLPs were pruned by setting a defined fraction of the weights to zero. Remaining non-zero weights were then re-trained with early stopping using the training and validation datasets, respectively. Finally, the Pearson correlation coefficient, mean squared error, and root mean square error was calculated and reported using the test dataset.

### Regression of deeply mutagenized sequences

Phenotypes and sequences or mutants were downloaded for WW domain [32, 33], eqFP611 [34], guanine-inhibited ribozyme Lib-2 [35], BRCA1 [36], and avGFP [37], with multiple sequence alignments generated *in silico*. For the guanine-inhibited ribozymes and BRCA1 RNA sequences, uracil (U) was replaced with thymine (T). Groups of identical sequences were assigned to training, validation, or test datasets as above. MLPs were then trained using the standard protMLP protocol, using the phenotypes as the target for regression.

### k-Nearest Neighbor regression

Phenotypes were predicted by k-Nearest Neighbor regression using the Hamming distance of the one-hot encoded sequence as the distance metric. The optimal number of neighbors was determined by systematically screening values between 1 and 20, selecting the k with the lowest MSE for the validation dataset.

### Random Forest regression

Phenotypes were predicted by random forest regression, testing forests of 10, 50, 100, 500, and 1000 trees and selecting the forest with the lowest MSE for the validation dataset. For each decision tree, the number of sequences required to split an internal node was limited to twice the number of trees in the forest to ensure overdetermination.

All calculations used custom scripts written in Python with the Biopython [38], Tensorflow [39], Keras [40], NumPy [41], SciPy [42], and Matplotlib [43] libraries.

## List of Abbreviations

ADK: Adenosine Kinase
CSP: Cold Shock Protein
kNN: k-Nearest Neighbor
MLP: Multi-layer perceptron
RF: Random Forest
SOD: Superoxide Dismutase

## Declarations

## Ethics approval and consent to participate

Not applicable.

## Consent for Publication

All authors provide their consent for publication.

## Availability of data and materials

The proposed methods are implemented in Python and publicly available at https://github.com/DavidBSauer/protMLP. The datasets used and/or analyzed during the current study are available from the corresponding author on reasonable request.

## Competing Interests

The authors declare that they have no competing interests

## Funding

This work was supported by the National Institutes of Health (R01-GM121994 and R01-NS108151 to D.N.W). D.B.S. was supported in part by a Postdoctoral Fellowship (PF-17-135-01) from the American Cancer Society and by the Office of the Assistant Secretary of Defence for Health Affairs, through the Peer Reviewed Cancer Research Program under Award No. W81XH-16-1-0153. Opinions, interpretations, conclusions, and recommendations are those of the authors and are not necessarily endorsed by the Department of Defence.

## Author Contributions

D.B.S. conceived of the study, designed the methodology, and analyzed the results. D.B.S and D.N.W. wrote the manuscript. D.N.W. supervised the research. All authors have approved the final version of the manuscript.

## Acknowledgements

The authors thank David Fenyo for helpful discussion of this work, and Jennifer Marden for critical review of this manuscript.

## Notes

https://github.com/DavidBSauer/protMLP

### Competing Interest Statement

The authors have declared no competing interest.

### Summary of Updates

Revisions to methods and analysis targets based on feedback.

