## Supplementary figures and tables for "Using machine learning to predict quantitative phenotypes from protein and nucleic acid sequences"

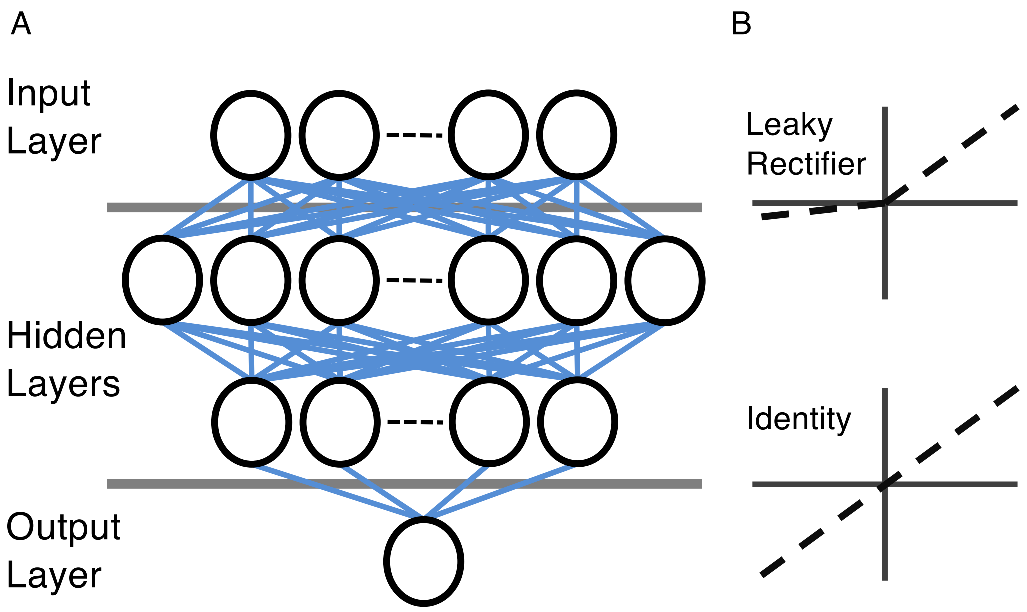


Figure S1. Design of a Multi-Layer Perceptron. (A) Structure of a MLP with 2 hidden layers. Connections between nodes shown as blue lines. Bias values are not shown. (B) Plots of Leaky Rectifier and Identity activation functions.

| | 1 | 2 | 3 |
|---|---|---|---|
| Seq1: | A | C | D |
| Seq2: | - | C | E |
| Seq3: | - | C | D |

⇩ One-hot encode sequences

| | A1 | -1 | C2 | D3 | E3 |
|---|---|---|---|---|---|
| Seq1: | 1 | 0 | 1 | 1 | 0 |
| Seq2: | 0 | 1 | 1 | 0 | 1 |
| Seq3: | 0 | 1 | 1 | 1 | 0 |

⇩ Remove invariant amino acids

| | A1 | -1 | D3 | E3 |
|---|---|---|---|---|
| Seq1: | 1 | 0 | 1 | 0 |
| Seq2: | 0 | 1 | 0 | 1 |
| Seq3: | 0 | 1 | 1 | 0 |

Figure S2. Encoding of protein sequence alignment. Aligned sequences are one-hot encoded, and invariant amino acids removed. Gap amino acids, representing a deletion in the sequence, are indicated by a dash mark.

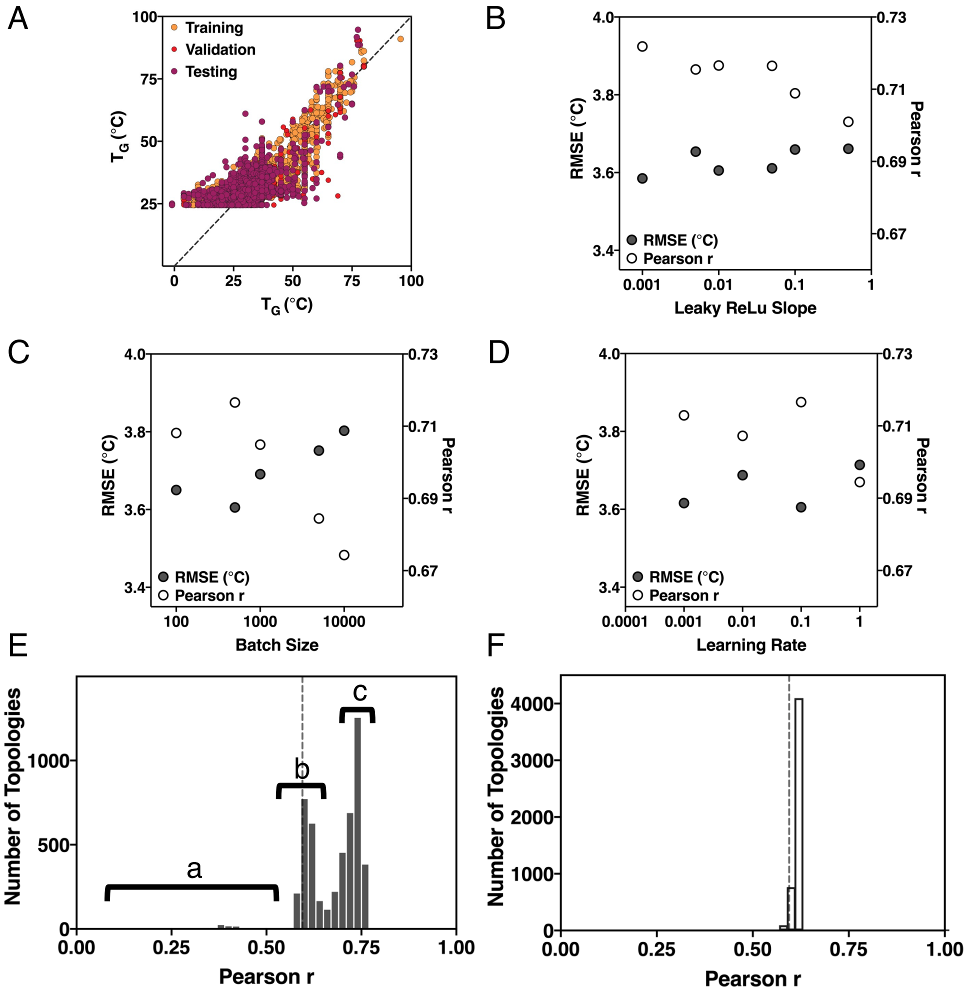


Figure S3. MLP accuracy is improved by optimization of the model hyperparameters. (A) Accuracy of MLP trained on Cold Shock Proteins from psychrophiles, mesophiles, and thermophiles, before optimization of model hyperparameters. Accuracy of MLPs in predicting organismal $T_G$ from CSP sequences while systematically scanning (B) the slope of the leaky ReLu activation function, (C) the training batch size, and (D) the learning rate. Accuracy in predicting the validation dataset for all trained MLPs, using either (E) a rectified or (F) identity activation function. The accuracy of a linear regression is indicated by the dotted line.

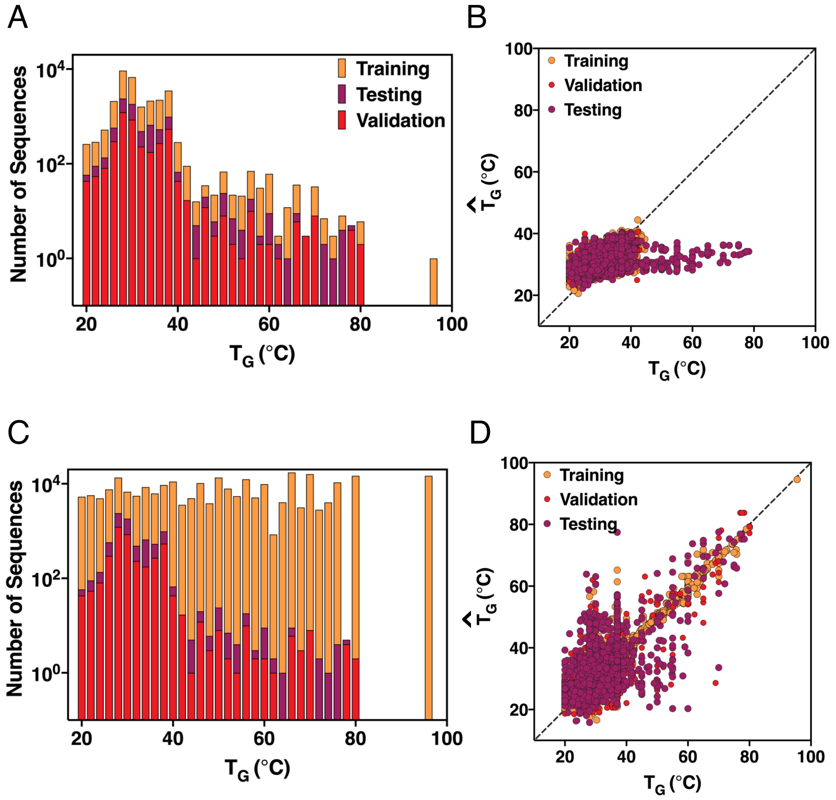


Figure S4. The CSP dataset is heavily skewed by mesophile sequences but can be balanced by over-sampling. (A) Log scale histogram of sequences by $T_G$. (B) Best rMLP regression using only mesophile training sequences. The dotted line indicates perfect prediction. (C) Distribution of sequence $T_G$ after balancing the training dataset by over-sampling. (D) Best rMLP using overbalanced training sequences. The dotted line indicates perfect prediction.

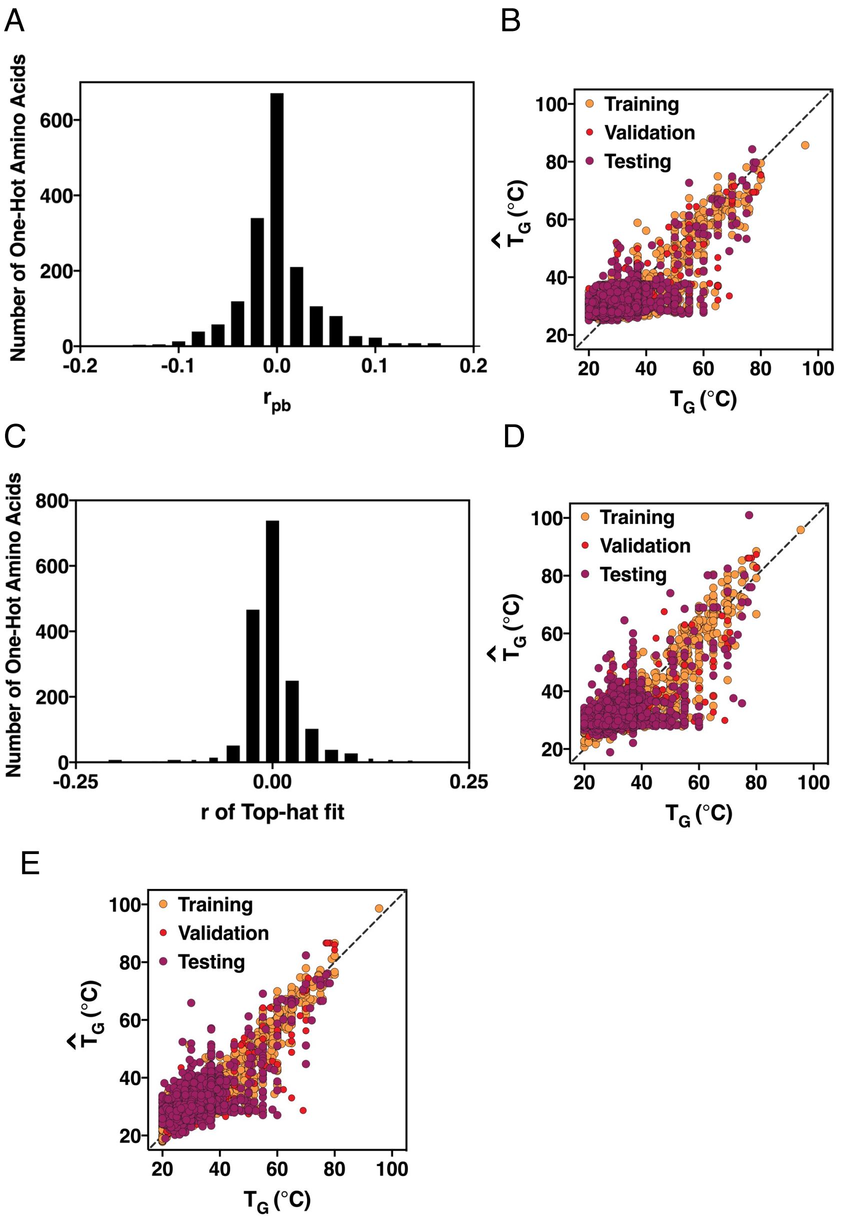


Figure S5. Only a subset of the training data is necessary. (A) First-order correlation of one-hot encoded amino acids to $T_G$. (B) Best rMLP regression using only one-hot encoded amino acids with an absolute $r_{pb}$ correlation greater than or equal to 0.1. The dotted line indicates perfect prediction. (C) Pearson correlation of a fit top-hat function to the one-hot encoded amino acids versus $T_G$. (D) Best rMLP regression using only one-hot encoded amino acids with an absolute top-hat correlation greater than or equal to 0.1. The dotted line indicates perfect prediction. (E) Pruned rMLP of underdetermined regression.

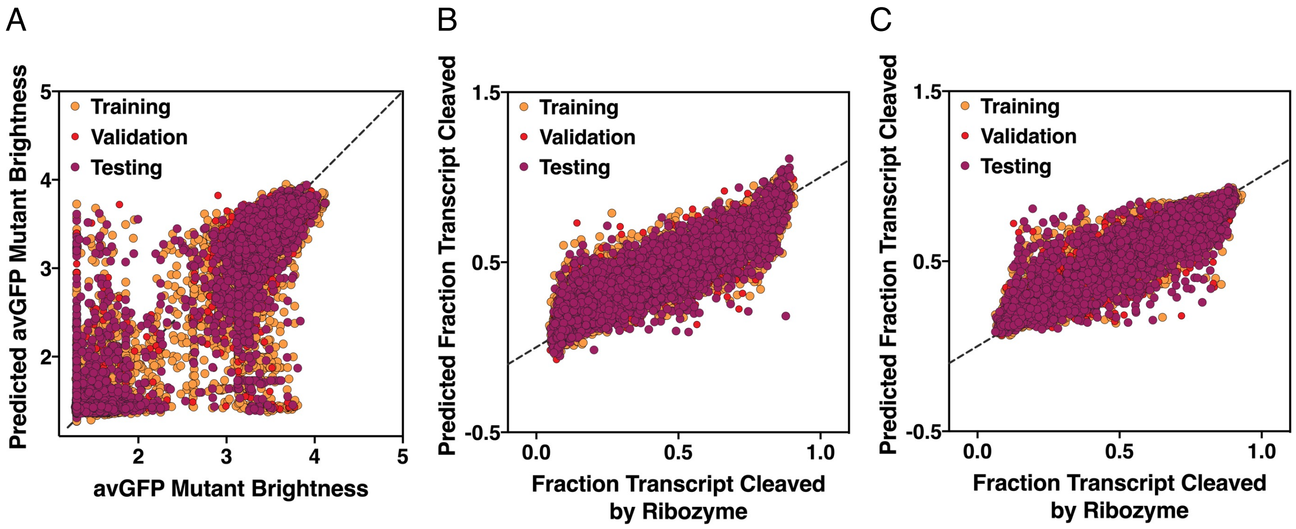


Figure S6. Predictions of deeply mutagenized protein and RNA sequences. (A) Regression of avGFP mutants' brightness. Regression of ribozyme activity (B) with and (C) without ligand. The dotted line indicates perfect prediction.

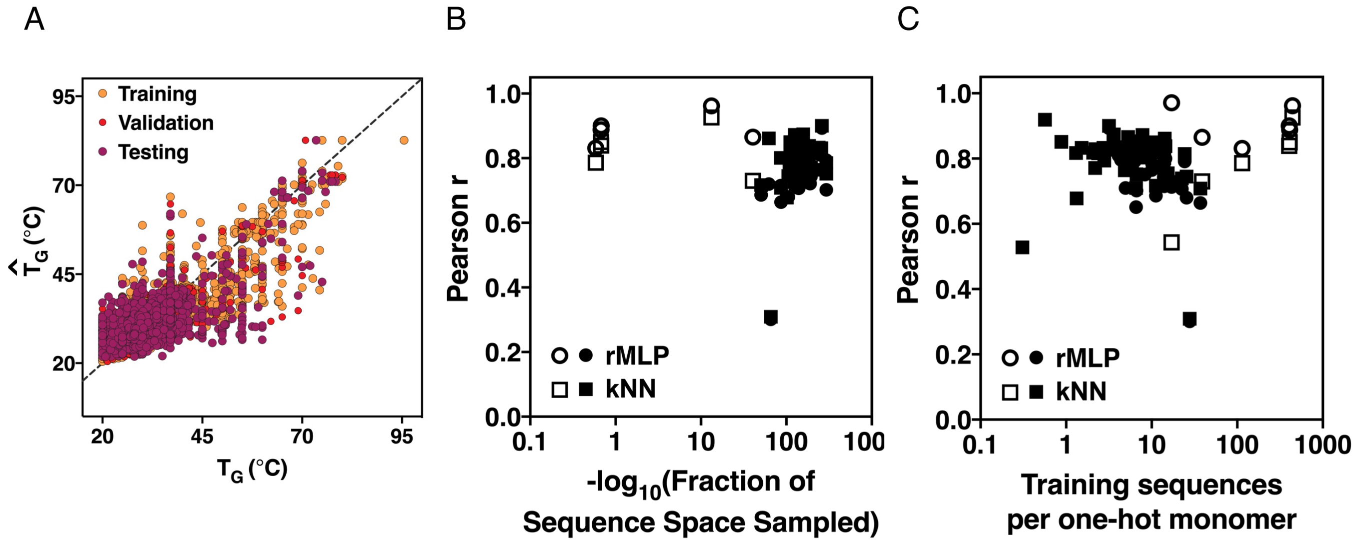


Figure S7. Regression methods which only consider sequence similarity can be used to predicted quantitative phenotypes. (A) kNN regression of CSP sequences. Dependence on rMLP and kNN accuracy on (A) number number of training sequences per one-hot amino acids or (B) fraction of sequence space sampled. Filled and empty shapes report on naturally-evolved and deep-mutagenesis regression, respectively.

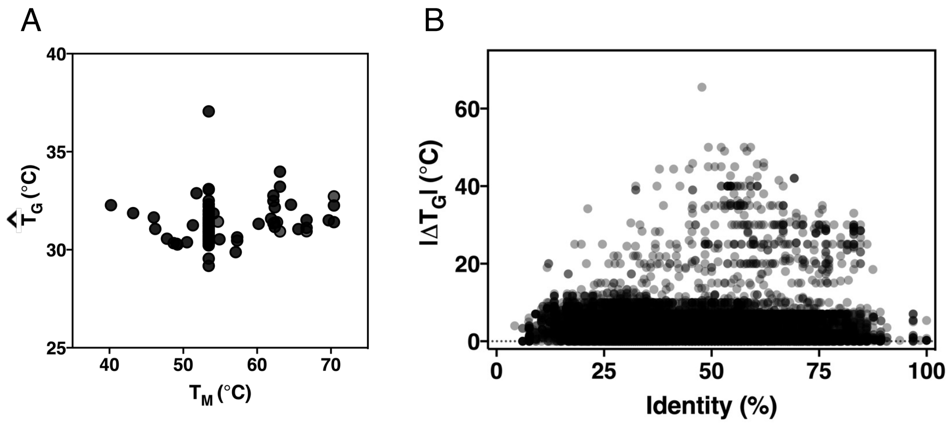


Figure S8. The CSP sequences contain insufficient depth for prediction of single and double mutants. (A) Predicted $T_G$ versus measured $T_M$ for BsCSP mutants. (B) Sequence identity to BsCSP versus absolute $T_G$ difference relative to *Bacillus subtillis* for all training data.

Table S1. Regression results

| | Pfam alignment | Phenotype | Sequences used | Pearson r Linear reg. | Pearson r rMLP reg. | Pearson r kNN reg. | Pearson r RFR reg. |
|---|---|---|---|---|---|---|---|
| Hsp20 family | PF00011 | $T_G$ | $T_G \geq 20$ | 0.731 | 0.808 | **0.818** | 0.780 |
| Cytochrome C family | PF00034 | $T_G$ | $T_G \geq 20$ | 0.571 | 0.702 | **0.752** | 0.682 |
| Globin family | PF00042 | $T_G$ | $T_G \geq 20$ | 0.825 | 0.826 | **0.874** | 0.860 |
| Cu/ZnSOD family | PF00080 | $T_G$ | $T_G \geq 20$ | 0.798 | | 0.833 | **0.841** |
| Fe/MnSOD N-terminal domain | PF00081 | $T_G$ | $T_G \geq 20$ | 0.745 | 0.827 | **0.872** | 0.871 |
| Thioredoxin family | PF00085 | $T_G$ | $T_G \geq 20$ | 0.680 | 0.800 | **0.838** | 0.795 |
| Tubulin/FtsZ family | PF00091 | $T_G$ | $T_G \geq 20$ | 0.810 | | 0.857 | **0.862** |
| Iron-Sulfur Cluster Binding domain | PF00111 | $T_G$ | $T_G \geq 20$ | 0.541 | 0.707 | **0.739** | 0.714 |
| ATP synthase subunit C | PF00137 | $T_G$ | $T_G \geq 20$ | 0.708 | 0.715 | 0.801 | **0.814** |
| Hsp10 family | PF00166 | $T_G$ | $T_G \geq 20$ | 0.771 | 0.814 | **0.831** | 0.812 |
| C2 domain | PF00168 | $T_G$ | $T_G \geq 20$ | 0.767 | 0.825 | 0.830 | **0.856** |
| 2-oxoacid Dehydrogenases Acyltransferase catalytic domain | PF00198 | $T_G$ | $T_G \geq 20$ | 0.776 | **0.828** | 0.820 | 0.807 |
| Cold Shock Protein | PF00313 | $T_G$ | $T_G \geq 20$ | 0.595 | 0.713 | **0.728** | 0.704 |
| Nucleoside diphosphate kinase | PF00334 | $T_G$ | $T_G \geq 20$ | 0.853 | 0.893 | **0.900** | 0.877 |
| Adenylate kinase | PF00406 | $T_G$ | $T_G \geq 20$ | 0.798 | | 0.818 | **0.846** |
| Chlorophyll A-B binding protein | PF00504 | $T_G$ | $T_G \geq 20$ | -0.059 | | 0.528 | **0.569** |
| Complex I domain | PF00662 | $T_G$ | $T_G \geq 20$ | 0.706 | 0.783 | **0.850** | 0.798 |
| OmpA outer membrane protein | PF00691 | $T_G$ | $T_G \geq 20$ | 0.602 | 0.721 | **0.755** | 0.713 |
| Flagellin C-terminal domain | PF00700 | $T_G$ | $T_G \geq 20$ | 0.735 | **0.825** | 0.818 | 0.780 |
| 3-hydroxyacyl-CoA dehydrogenase C-terminal domain | PF00725 | $T_G$ | $T_G \geq 20$ | 0.676 | 0.776 | **0.823** | 0.785 |
| GrpE nucleotide exchange factor | PF01025 | $T_G$ | $T_G \geq 20$ | 0.794 | | **0.829** | 0.764 |

| | | | | | | | |
|---|---|---|---|---|---|---|---|
| MarR repressor | PF01047 | $T_G$ | $T_G \geq 20$ | 0.480 | 0.664 | **0.708** | 0.692 |
| Omp85 outer membrane protein | PF01103 | $T_G$ | $T_G \geq 20$ | 0.814 | | 0.851 | **0.881** |
| Prokaryote Diacylglycerol Kinase | PF01219 | $T_G$ | $T_G \geq 20$ | 0.746 | 0.796 | **0.835** | 0.829 |
| Prokaryote Cytochrome b561 | PF01292 | $T_G$ | $T_G \geq 20$ | 0.758 | 0.799 | 0.794 | **0.818** |
| Exonuclease domain | PF01367 | $T_G$ | $T_G \geq 20$ | 0.726 | 0.817 | **0.853** | 0.791 |
| Cytochrome C assembly protein | PF01578 | $T_G$ | $T_G \geq 20$ | 0.765 | | **0.816** | 0.792 |
| Peptide Methionine Sulfoxide Reductase | PF01625 | $T_G$ | $T_G \geq 20$ | 0.760 | **0.814** | 0.764 | 0.790 |
| Sodium/Calcium exchanger | PF01699 | $T_G$ | $T_G \geq 20$ | 0.750 | 0.845 | **0.866** | 0.828 |
| Large-conductance mechanosensitive channel | PF01741 | $T_G$ | $T_G \geq 20$ | 0.758 | | **0.833** | 0.786 |
| MthK channel regulatory C-terminal domain | PF02080 | $T_G$ | $T_G \geq 20$ | 0.607 | 0.751 | **0.803** | 0.777 |
| MthK channel regulatory N-terminal domain | PF02254 | $T_G$ | $T_G \geq 20$ | 0.661 | 0.765 | **0.819** | 0.750 |
| MlaD lipid transport protein | PF02470 | $T_G$ | $T_G \geq 20$ | 0.660 | 0.743 | **0.772** | 0.764 |
| Fluc channel | PF02537 | $T_G$ | $T_G \geq 20$ | 0.719 | 0.809 | **0.831** | 0.792 |
| 3-hydroxyacyl-CoA dehydrogenase NAD-binding domain | PF02737 | $T_G$ | $T_G \geq 20$ | 0.705 | **0.794** | 0.785 | 0.764 |
| Cache domain | PF02743 | $T_G$ | $T_G \geq 20$ | 0.709 | | **0.771** | 0.731 |
| Fe/Mn SOD C-terminal domain | PF02777 | $T_G$ | $T_G \geq 20$ | 0.729 | 0.813 | **0.831** | 0.803 |
| Thioesterase family | PF03061 | $T_G$ | $T_G \geq 20$ | 0.597 | 0.791 | **0.814** | 0.775 |
| 3D domain | PF06725 | $T_G$ | $T_G \geq 20$ | 0.701 | **0.786** | 0.775 | 0.781 |
| Two-component Histidine Kinase domain | PF07730 | $T_G$ | $T_G \geq 20$ | 0.557 | 0.680 | **0.745** | 0.716 |
| Potassium Channel | PF07885 | $T_G$ | $T_G \geq 20$ | 0.676 | 0.792 | 0.820 | **0.827** |
| 1-Cys Peroxiredoxin C-terminal domain | PF10417 | $T_G$ | $T_G \geq 20$ | 0.747 | 0.720 | **0.861** | 0.813 |
| FtsZ C-terminal domain | PF12327 | $T_G$ | $T_G \geq 20$ | 0.791 | **0.836** | 0.797 | 0.831 |

| | | | | | | | |
|---|---|---|---|---|---|---|---|
| Nicotinamide Nucleotide Transhydrogenase | PF12769 | $T_G$ | $T_G \geq 20$ | 0.720 | 0.710 | 0.783 | **0.805** |
| ECF transporter S-subunit | PF12822 | $T_G$ | $T_G \geq 20$ | 0.766 | | **0.817** | 0.747 |
| PPR repeat | PF13041 | $T_G$ | $T_G \geq 20$ | 0.209 | 0.301 | 0.309 | **0.500** |
| GAF domain | PF13185 | $T_G$ | $T_G \geq 20$ | 0.570 | 0.651 | **0.710** | 0.683 |
| Cytochrome B | PF13631 | $T_G$ | $T_G \geq 20$ | 0.087 | | **0.919** | 0.914 |
| Keratin domain | PF13885 | $T_G$ | $T_G \geq 20$ | 0.618 | | **0.678** | 0.589 |
| 2-oxoglutarate dehydrogenase N-terminal domain | PF16078 | $T_G$ | $T_G \geq 20$ | 0.671 | 0.686 | **0.717** | 0.708 |
| eqFP611 | deep mutagenesis | Brightness | all | 0.351 | **0.962** | 0.926 | 0.959 |
| WW domain | deep mutagenesis | Binding Affinity | all | 0.742 | **0.865** | 0.730 | 0.715 |
| Ribozyme | deep mutagenesis | Activity | Gln+ | 0.496 | **0.888** | 0.850 | 0.853 |
| Ribozyme | deep mutagenesis | Activity | Gln- | 0.503 | **0.900** | 0.839 | 0.841 |
| BRCA1 | deep mutagenesis | Enrichment | all | 0.579 | **0.831** | 0.786 | 0.799 |
| avGFP | deep mutagenesis | Brightness | all | 0.829 | **0.971** | 0.543 | 0.861 |
| 3-hydroxyacyl-CoA dehydrogenase C-terminal domain | PF02470 | pH | all | 0.598 | | 0.611 | **0.622** |
| YceI periplasmic protein | PF04264 | pH | all | 0.426 | | 0.618 | **0.665** |

| | | | | Linear AUC | rMLP AUC | kNN AUC | RFR AUC |
|---|---|---|---|---|---|---|---|
| Fe/Mn SOD N-terminal domain | PF00081 | Oxygen Tolerance | Aerobe or Anaerobe | 0.974 | **0.976** | 0.959 | 0.968 |
| Fe/Mn SOD C-terminal domain | PF02777 | Oxygen Tolerance | Aerobe or Anaerobe | 0.962 | 0.965 | 0.957 | **0.979** |
